# A variant in *TAF1* is associated with a new syndrome with severe intellectual disability and characteristic dysmorphic features

**DOI:** 10.1101/014050

**Authors:** Jason Ou’Rawe, Yiyang Wu, Alan Rope, Laura T. Jimenez Barrón, Jeffrey Swensen, Han Fang, David Mittelman, Gareth Highnam, Reid Robison, Edward Yang, Kai Wang, Gholson Lyon

## Abstract

We describe the discovery of a new genetic syndrome, RykDax syndrome, driven by a whole genome sequencing (WGS) study of one family from Utah with two affected male brothers, presenting with severe intellectual disability (ID), a characteristic intergluteal crease, and very distinctive facial features including a broad, upturned nose, sagging cheeks, downward sloping palpebral fissures, prominent periorbital ridges, deep-set eyes, relative hypertelorism, thin upper lip, a high-arched palate, prominent ears with thickened helices, and a pointed chin. This Caucasian family was recruited from Utah, USA. Illumina-based WGS was performed on 10 members of this family, with additional Complete Genomics-based WGS performed on the nuclear portion of the family (mother, father and the two affected males). Using WGS datasets from 10 members of this family, we can increase the reliability of the biological inferences with an integrative bioinformatic pipeline. In combination with insights from clinical evaluations and medical diagnostic analyses, these DNA sequencing data were used in the study of three plausible genetic disease models that might uncover genetic contribution to the syndrome. We found a 2 to 5-fold difference in the number of variants detected as being relevant for various disease models when using different sets of sequencing data and analysis pipelines. We derived greater accuracy when more pipelines were used in conjunction with data encompassing a larger portion of the family, with the number of putative de-novo mutations being reduced by 80%, due to false negative calls in the parents. The boys carry a maternally inherited missense variant in a X-chromosomal gene *TAF1*, which we consider as disease relevant. TAF1 is the largest subunit of the general transcription factor IID (TFIID) multi-protein complex, and our results implicate mutations in *TAF1* as playing a critical role in the development of this new intellectual disability syndrome.

## 1 Introduction

Dramatic cost reductions and rapid advancements in the development of efficient sequencing technologies [1–3] have led to widespread use of exome and whole genome sequencing (WGS) [4–6] in a variety of research and clinical settings [7]. Computational tools for processing and analyzing these sequence data have been developed in parallel, and many are now freely available and straightforward to implement [4]. In this context, biomedical exome sequencing and WGS has led to the discovery of the genetic basis for many conditions, including Miller Syndrome [8] and others [9].

The intellectual impetus for this study included how to identify the major genetic contribution to a particular syndrome in only one proband or two affected siblings, in the absence of other affected people with the same known syndrome. This task is much easier in the presence of multiple affected people spread out over two or more generations [10, 11], but this is usually not the case for most biomedical presentations of idiopathic disorders. Genetic discovery can also be easier in consanguineous pedigrees with autosomal recessive conditions [12, 13], but such consanguinity is not widespread in many parts of the world [14]. We instead focused on an analysis that initially included one family with only two siblings that are both afflicted by an idiopathic syndrome, and so we utilized WGS to cover as much of the genome as we could, particularly as any number of mutated nucleotides in the genome might influence some phenotype during embryogenesis or postnatal life [15–33].

This Caucasian family was recruited from Utah, USA. Illumina-based WGS was performed on 10 members of this family, with additional Complete Genomics-based WGS performed on the nuclear portion of the family (mother, father and the two affected males). Sequence data were processed with a number of bioinformatics pipelines. Using comprehensive datasets generated by an aggregation of results stemming from these pipelines, we can increase the reliability of the biological inferences stemming from these data. In combination with insights from clinical evaluations and medical diagnostic analyses, these DNA sequencing data were used in the study of three plausible genetic disease models that might uncover genetic contribution to the syndrome.

## 2 Materials and Methods

This study consists of two methodological components, clinical and genomic sequencing/analysis. The clinical component includes research participant enrollment, clinical evaluation, diagnostic analyses and a detailed clinical report. The genomic sequencing component includes whole genome sequencing as well as downstream analyses aimed at annotating and assigning biological function to detected genomic variants.

### A. Clinical methods

#### A.1. Enrollment of research participants

The collection and the analysis of DNA was conducted by the Utah Foundation for Biomedical Research, as approved by the Institutional Review Board (IRB) (Plantation, Florida). Written informed consent was also obtained from all study participants, and research was carried out in compliance with the Helsinki Declaration.

#### A.2. Clinical evaluation/diagnostics

A broad range of clinical diagnostic testing was performed on both affected male siblings, including karyotyping, a high resolution X-chromosome CGH array (720K Chromosome X Specific Array from Roche NimbleGen, Inc. USA), subtelomeric FISH study, methylation study for Angelman syndrome, *XNP* sequencing for ATRX, and fragile X DNA testing. In addition, we performed diagnostics on serum amino acid levels, urine organic acids levels, sweat chloride levels, plasma carnitine profile, and immunoglobulin levels. We also performed urine mucopolysaccharidosis (MPS) screening and examined thyroid profiles. Cranial ultrasound was performed and brain imagery was obtained using magnetic resonance imaging (MRI) and computed tomography (CT) scanning techniques. Images of the spine were also obtained using MRI. Moreover, cerebrospinal fluid (CSF) was collected from the elder sibling, and neurotransmitter metabolites, tetrahydrobiopterin (BH4) and neopterin (N) profile were screened.

#### A.3. Custom *X* CGH array and *X*-chromosome skewing assay

500 ng of each research participant’s DNA was labeled with 5’-Cy3 tagged nanomers (NimbleGen) while a female control was labeled with Cy5 nonamers. After purification by isopropanol precipitation, 31 ug each of labeled research participant and reference DNA were combined. The mixture was hybridized to a custom NimbleGen 720K Chromosome X Specific Array for 42 hours at 42C in a MAUI Hybridization System (BioMicro Systems). The array was then washed according to the manufacturer’s recommendation (Nimblegen) and immediately scanned. After scanning, fluorescence intensity raw data was extracted from the scanned images of the array using NimbleScan v2.6 software. For each of the spots on the array, normalized log2 ratios of the Cy3-labeled research participant sample vs the Cy5-reference sample were generated using the SegMNT program. The data was visualized with Nexus 6.1 software (Biodiscovery).

X-chromosome skewing assay analyses were performed using an adaptation of the technique described by Allen et al. (1992) [34].

### B. Whole Genome Sequencing and analysis methods

Two different sequencing strategies were employed. Initial sequencing efforts focused on whole genome sequencing using the Complete Genomics (CG) sequencing and analysis pipeline v2.0 for the mother, father and two affected boys. Additional whole genome sequencing was performed subsequent to this initial effort, using the Illumina HiSeq 2000 sequencing platform. The mother, father, two affected boys and six other immediate family members were sequenced using the Illumina HiSeq 2000. Raw sequencing data stemming from Illumina sequencing was processed by a variety of analysis pipelines and subsequently pooled with variants detected by the CG sequencing an analysis pipeline. Downstream and functional annotation tools were used to evaluate variants detected by the various methods.

#### B.1. Complete Genomics whole genome sequencing and variant detection

After quality control to ensure lack of genomic degradation, we sent 10ug DNA samples to Complete Genomics (CG) at Mountain View, California for sequencing. The whole-genome DNA was sequenced with a nanoarray-based short-read sequencing-by-ligation technology, including an adaptation of the pairwise end-sequencing strategy. Reads were mapped to the Genome Reference Consortium assembly GRCh37. Due to the proprietary data formats, all the sequencing data QC, alignment and variant calling were performed by CG as part of their sequencing service, using their version 2.0 pipeline.

#### B.2. Illumina HiSeq 2000 whole genome sequencing and variant detection

After the samples were quantified using Qubit dsDNA BR Assay Kit (Invitrogen), 1ug of each sample was sent out for whole genome sequencing using the Illumina® Hiseq 2000 platform. Sequencing libraries were generated from 100ng of genomic DNA using the Illumina TruSeq Nano LT kit, according to manufacturer recommendations. The quality of each library was evaluated with the Agilent bioanalyzer high sensitivity assay (less than 5% primer dimers), and quantified by qPCR (Kappa Biosystem, CT). The pooled library was sequenced in three lanes of a HiSeq2000 paired end 100bp flow cell. The number of clusters passing initial filtering was above 80%, and the number of bases at or above Q30 was above 85%.

Illumina reads were mapped to the hg19 reference genome using BWA v0.6.2-r126, and variant detection was performed using the GATK v. 2.8-1-g932cd3a. A second analytical pipeline was used to map the Illumina reads and detect variants using novoalign v3.00.04 and the FreeBayes caller v9.9.2-43-ga97dbf8. Additional variant discovery procedures included Scalpel v0.1.1 for insertion or deletion (INDEL) detection, RepeatSeq v0.8.2 for variant detection in short tandem repeat regions, and the ERDS (estimation by read depth) method v1.06.04 and PennCNV (2011Jun16 version) for detecting larger copy number variants (CNVs).

We used several methods to prioritize and identify possible disease-contributory germ-line variants, including VAAST [10, 35], Golden Helix SVS v8.1.4 [36], ANNOVAR (2013Aug23 version) [37], and GEMINI v0.9.1 [38]. VAAST employs a likelihood-based statistical framework for identifying the most likely disease-contributory variants given genomic makeup and population specific genomic information. SVS, ANNOVAR and GEMINI employ more traditional annotation and filtering-based techniques that leverage data stored in public genomic databases (i.e., dbSNP 137, 1000 Genomes phase 1 data, NHLBI 6555 exomes, etc.).

We used two distinct variant prioritization schemes. The first scheme, which we will refer to as the ‘coding’ scheme, requires all variants to be within a coding region of the genome. Splice site variants are also included. The second scheme, which we will refer to as the ‘CADD’ scheme, requires all variants to have a CADD score of >20. CADD scores do not preclude non-coding genetic variants from the resulting list of potentially deleterious variants. Both schemes required each variant to have a low population frequency (MAF < 1%). Prioritized variants were manually verified by inspecting sequence alignments using Golden Helix GenomeBrowse v2.0.3 [http://www.goldenhelix.com/GenomeBrowse/index.html]. A signal was considered plausible if 4 or more reads supported the alternative allele in more than one family member. Population frequency information was corroborated using the NCBI Variation viewer [http://www.ncbi.nlm.nih.gov/variation/view/]. See the Supplementary Information for details about the variant calling and prioritization analyses.

## 3 Results

### A. Clinical evaluations and phenotypic presentation

The initial probands selected for study by the corresponding author (GJL) were two affected brothers, ages 12-and 14-years-old respectively, with severe ID, autistic behaviors, anxiety, attention deficit hyperactivity disorder, and very distinctive facial features. Among the facial features are a broad, upturned nose, sagging cheeks, downward sloping palpebral fissures, prominent periorbital ridges, deep-set eyes, relative hypertelorism, thin upper lip, a high-arched palate, prominent ears with thickened helices, and a pointed chin. Other shared phenotypic symptoms include strabismus (exotropia), blocked tear ducts, microcephaly, mild ventriculomegaly, deficiency of the septum pellucidum, hypoplasia of the corpus callosum, low cerebral white matter volume, oculomotor dysfunction, frequent otitis media with effusion, hearing impairments (mixed conductive/sensorineural), oral motor dysphagia, kyphosis, a peculiar gluteal crease with sacral caudal remnant (without any spinal abnormalities), dysplastic toenails, hyperextensible joints (especially fingers and wrists), spasticity, ataxia, gait abnormalities, growth retardation and global developmental delays, especially in the areas of gross motor and verbal expression. The younger of the two affected siblings also suffers from frequent episodes of contact dermatitis and eczema, scoliosis, sleep-wake dysregulation, as well as asthma, although he no longer requires medication for the latter. The elder brother, on the other hand, has diplegia, and has received Botulinum Toxin (Botox) therapy for his lower-extremity spasticity for six years. A review of systems (ROS) questionnaire revealed no other obvious, shared or otherwise, symptoms or malformations. See Table 1 for a summary of the clinical features of the affected male siblings.

**Tabel 1.**
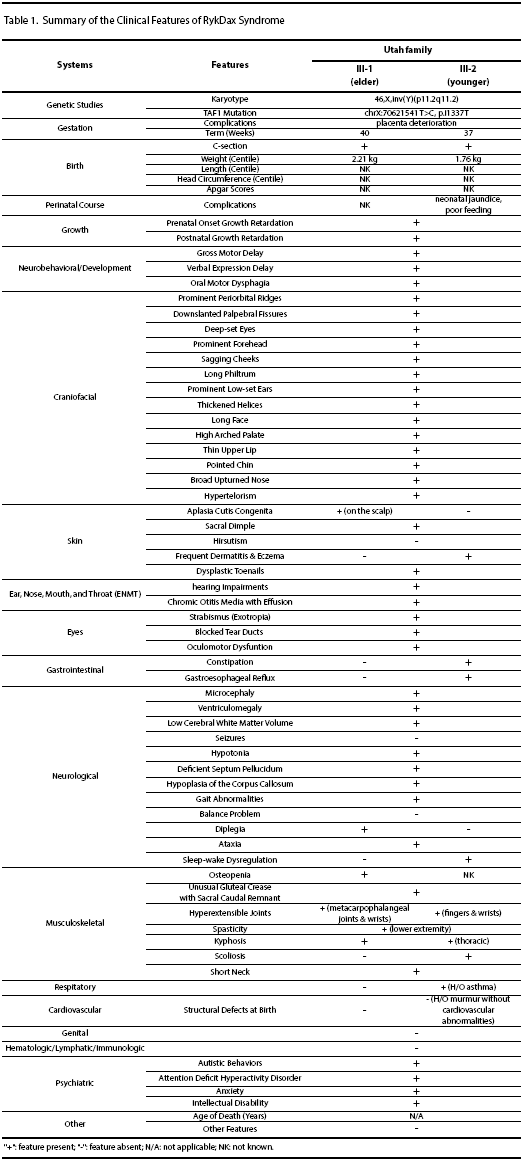
This table illustrates known clinical features across the affected individuals, as well as other noted clinical characteristics on these individuals.

The parents of the two affected siblings are non-consanguineous and are both healthy. The mother has been evaluated for PKU and had normal plasma amino acid levels. The family history does not reveal any members, living or deceased, with phenotypic or syndromic characteristics that resemble the described syndrome, and there is a male cousin who is unaffected. An X-chromosome skewing assay revealed that the mother of the two affected boys has skewed, 99:1, X-chromosome inactivation. The grandmother, as well as the aunt of the affected boys, does not show any appreciable X-chromosome skewing (Figure 1), which suggested the possibility of a newly arising deleterious X-chromosome variant, although it is well established that this also could be non-specific [39].

**Figure 1.**
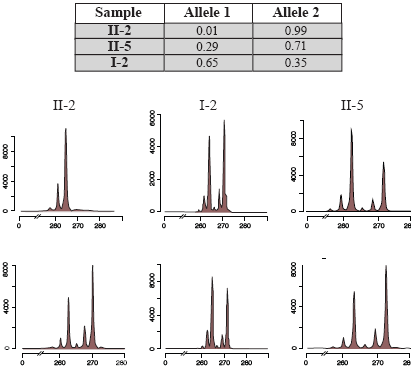
X-chromosome inactivation assays were performed on the mother (**II-2**), aunt (**II-5**), and maternal grandmother (**I-2**) of the two affected siblings. The assay reveals normal X-chromosome inactivation ratios in the aunt and maternal grandmother, however the mother of the affected siblings exhibits a 99:1 X-chromosome inactivation ratio.

Both pregnancies with these male fetuses were complicated by placenta deterioration, and both affected siblings were diagnosed with intra-uterine growth retardation (IUGR) and were eventually delivered through Caesarean section (C-section). The mother denied any alcohol or drug use, nor any exposure to environmental toxins during the course of both pregnancies. The elder boy was born in the 40th gestational week with a birth weight of 2.21 kg and a notable birth defect of aplasia cutis congenita, which was surgically corrected at the age of 4 days old. The younger boy was born in the 37th gestational week with a birth weight of 1.76 kg. A heart murmur was noticed at his birth, but echocardiography confirmed the absence of any further or more serious cardiovascular abnormalities. He was treated with light for neonatal jaundice, and required a feeding tube during the first few days of his life due to difficulties swallowing and digesting food. During the most recent examinations, the younger boy (aged 1011/12 years) had a height of 129.7 cm (2% tile), a weight of 30.8 kg (19% tile, BMI 18.3 kg/m2), and his occipital frontal circumference (OFC) was 51 cm (4.5th percentile); while his elder brother (aged 1111/12 years) had a height of 136.8 cm (5% tile), a weight of 26.3 kg (0% tile, BMI 14.1 kg/m2), and his OFC was 49.5 cm (0.2th percentile) at the time.

Brain MRIs of the two brothers demonstrated a remarkably similar constellation of abnormalities. In both subjects, there was hypoplasia of the isthmus and splenium of the corpus callosum with thickness falling below the third percentile reported for individuals of the same age [40]. As is often the case with callosal hypoplasia, there was associated dysmorphic configuration of the lateral ventricles and mild lateral ventriculomegaly without positive findings of abnormal CSF dynamics (i.e. no imaging evidence of hydrocephalus). There was also deficiency of the septum pellucidum in both brothers, with the older brother having absence of the posterior two-thirds of the septum pellucidum and the younger brother having complete absence of the septal leaflets. Findings associated with septooptic dysplasia included underdeveloped pituitary glands for age, deficiency of the anterior falx with mild hemispheric interdigitiation, and question of small olfactory bulbs despite fully formed olfactory sulci. However, the optic nerves appeared grossly normal in size. Finally, there was subjective vermian hypoplasia with the inferior vermis resting at the level of the pontomedullary junction rather than a more typical lower half of the medulla. Pertinent negatives included absence of a malformation of cortical development, evidence of prior injury, or conventional imaging evidence of a metabolic/neurodegenerative process.

Other clinical diagnostic testing performed on both affected siblings (see Clinical Methods section) did not reveal any known disorders. Although chromosomal analysis revealed that both boys have the karyotype of 46,X,inv(Y)(p11.2q11.2), this is known to be a normal population variant.

### B. Whole genome sequencing and bioinformatics analysis

#### B.1. Whole genome sequencing

With our effort to move beyond exome sequencing and into whole genome analyses, over the past few years we have developed and published comprehensive whole genome analysis pipelines, including for finding insertions and deletions (INDELs) [7, 41–46]. We and others have shown that the two dominant WGS platforms (Illumina and Complete Genomics) are complementary, as both miss variants [42, 47], and so both platforms were used in this study. Initial sequencing efforts focused on whole genome sequencing using the Complete Genomics (CG) sequencing and analysis pipeline v2.0 for the mother, father and two affected boys. Additional whole genome sequencing was performed subsequent to this initial effort, using the Illumina HiSeq 2000 sequencing platform on the mother, father, two affected boys and six other immediate family members. Raw sequencing data stemming from Illumina sequencing was processed by a variety of analysis pipelines and subsequently pooled with variants detected by the CG sequencing and analysis pipeline (see Whole Genome Sequencing and Analysis Methods section). Downstream and functional annotation tools were used to evaluate variants detected by the various methods. Complete Genomics WGS was optimized to cover 90% of the exome with 20 or more reads and 85% of the genome with 20 or more reads. Illumina WGS resulted in an average mapped read depth coverage of 37.8X (SD=1.3X). >90% of the genome was covered by 30 reads or more and >80% of the bases had a quality score of >30. See Supplementary Table 1 for more details about the sequencing data.

#### B.2. Bioinformatics analyses and variant calling

The mean number of variants per individual that were detected using the Illumina sequencing data across all of the detection methods was 3,583,905.1 (SD=192,317.5) SNPs, 650,708.2 (SD=84,125.7) INDELs (6,310.2 mean INDELs from Scalpel with a SD of 74.3, as it only detects signals in exon regions), 1,338,503 (SD=7,622.2) variants observed in STR regions, and 327.4 CNVs with a SD of 8.2 (and a mean of 49.1 and a SD of 15.8 for CNVs from PennCNV, which detects signals over a smaller search space than ERDS). For the CG sequence data, the mean number variants per individual detected are 3,457,584 (SD=51,665.8), 565,691.5 (SD=16,247.7), and 175.3 (SD=12.8) for SNPs INDELs and CNVs respectively. 14 unique INDELS and SNVs were discovered using two different prioritization schemes, with only a single coding SNV being reliably identified by both schemes. No known disease-contributory CNVs were discovered, but we archive in our study 8 de-novo CNVs that are not currently associated with any biological phenotype (see Supplementary Table 2 for the list of CNVs).

**Table 2.**
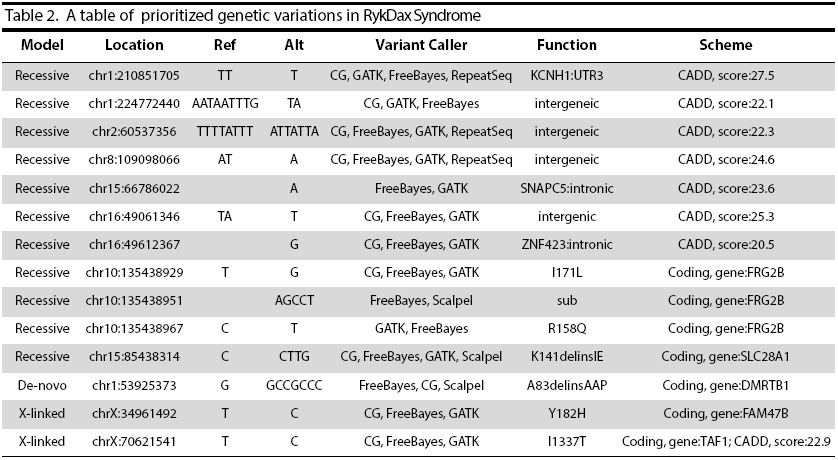
Variants conforming to the three disease models, de-novo, autosomal recessive and X-linked were identified. We show a list resulting from the CADD prioritization scheme as well as from the coding prioritization scheme. Both schemes required each variant to have a low population frequency (MAF < 1%). The coding scheme required all variants to also be within a coding region of the genome and to be a non-synonymous change. The CADD scheme requires all variants to have a CADD score of >20, along with the aforementioned population frequency. A variation in *TAF1* was the only variation to be reliably detected using both prioritization schemes.

#### B.3. Variant prioritization

Using the two variant prioritization schemes described in the Materials and Methods section, we discovered a set of putative variants from among the three disease models tested here. We found 7 potentially important variants in non-coding regions of the genome and 7 in coding regions across the two prioritization schemes (Table 2). These variants fall within coding and non-coding regions of a total of 8 known genes, *TAF1*, *FAM47B*, *SLC28A1*, *FRG2B*, *DMRTB1*, *ZNF423*, *SNAPC5* and *KCHN1*. We found a number of CNVs that were not known to be associated with any deleterious phenotype that we could find (Supplementary Table 2). As stated above, only one variant was shared between the two different schemes employed here, namely a non-synonymous change in *TAF1* that resulted in an isoleucine (hydrophobic) to threonine (polar) change on the 1337th residue. The protein change occurs within a linker region before two bromodomains and after a co-factor (*TAF7*) interacting domain. This linker region is highly conserved in multicellular eukaryotes (Figure 2B), but is not present in S. cerevisiae TAF1. Therefore, this linker region is not located in a recently reported crystal structure for the S. cerevisiae TAF1-TAF7 complex [49].

**Figure 2.**
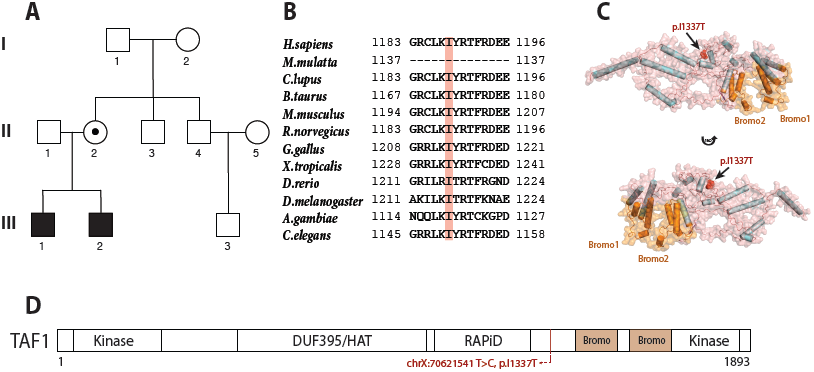
(**A**) Pedigree drawings of the Utah family, (**B**) A protein sequence alignment of TAF1 between *H.sapiens*, *M.mulatta*, *C.lupus*, *B.taurus*, *M.musculus*, *R.norvegicus*, *G.gallus*, *X.tropicalis*, *D.rerio*, *D.melanogaster*, *A.gambiae, and C.elegans*, listed from top to bottom. This alignment was generated using the MUSCLE[48] software in the HomoloGen [https://www.ncbi.nlm.nih.gov/homologene] website. The TAF1 I1337T location is highlighted in red, which shows a high degree of protein sequence conservation, (**C**) computational modeling of the TAF1 protein structure from residues 1080 to 1579 using I-TASSER and (**D**) known *TAF1* domains, with the *TAF1* variant indicated.

#### B.4. Variant scores

The *TAF1* variant found in this family was ranked highest among the variants being tested with VAAST using an X-linked model (with a p-value of 0.00184 and a rank of 14.59) and it is ranked highly in terms of its potential functional significance. This variant is ranked by CADD as being within the top l% most delete-rious variants in the human genome, it is scored by PolyPhen-2 [50] as being “Probably Damaging” with a score of 0.996, by SIFT [51] as “Damaging” with a score of 0.003, and also by PROVEAN [52] as “Deleterious” with a score of -3.51. This variant in *TAF1* is novel, as it is not found in public databases (i.e., dbSNP 137, 1000 Genomes phase 1 data, NHLBI 6500 exomes, or ExAC version 0.2).

#### B.5. Protein modeling

To investigate how the *TAF1* variant may influence protein structure and packaging, we built a structure model for the region of residues 1080 to 1569 using I-TASSER (Figure 2C). Residues spanning 1120 to 1270 contain the RAP74 interacting do-main (RAPiD), which has been shown to be important for the interactions between TAFl and RAP74 (GTF2F1) and TAF1-TAF7 [53]. Residues spanning 1373 to 1590 contain two Bromo domains (Bromo1: 1397-1467 and Bromo2: 1520-1590, Figure 2C and 2D). These Bromo do-mains consist of a bundle of four alpha helices that form a hydrophobic pocket to recognize an acetyl lysine, such as those on the N-terminal tails of histones. The variant found in this family, I337T, is located between the RAPiD and Bromo domains (shown as red spheres). Although this variant is not within any known protein domain, we speculate that it may affect domain packing of TAF1, which may interfere with the TAF1-TAF7 interacting surface [49, 53] or mark the protein for proteolytic degradation.

## Discussion

In general, we found benefits in using multiple informatics pipelines with WGS across a multi-generational pedigree for the identification and prioritization of human genetic variation potentially important in the disease phenotype discussed. We highlight here a variant found on the X-chromosome of the two affected male boys from Utah, USA, which was transmitted to them from their mother who acquired this variant spontaneously (de novo) and who is herself affected by extreme X-chromosome skewing. We were able to prioritize variants that were not observed in other members of the sequenced family, including not being present in an unaffected male cousin. The only variant found in both of our two variant prioritization schemes is in a highly conserved region of TAF1, which is the largest subunit of the TFIID multi-protein complex involved in transcription initiation [49, 54-57].

### A. *TAF1* variant implicated in disease phenotype

TAF1 (TATA-box-binding protein associated factor 1) is part of the TFIID multi-protein complex, which consists of the TATA binding protein (TBP) and 12 additional TBP associated factors (TAFs). TFIID has been implicated in promoting transcription initiation by recognizing promoter DNA and facilitating the nucleation of other factors to aid in the assembly of the pre-initiation complex [55]. TFIID also interacts with transcriptional activators as a co-activator [55]. TAF1 is the largest known TFIID associated TAF, and it binds directly to the TBP via a conserved N-terminal domain. Through its binding to TBP, TAF1 is thought to influence some control over the activation of genes promoted by TATA or other DNA motifs by inhibiting the TBP subunit from binding to these regions [54]. More recent work has reported aberrant expression affecting hundreds of *D. melanogaster* genes as a result of *TAF1* transcript depletion, with many more of these genes being expressed more vigorously while a smaller number are expressed less vigorously than under wild-type conditions [58]. In addition and consistent with the notion that TAF1 is important in controlling the binding patterns of TFIID to specific promoter regions, this study showed that the set of genes conferring increased expression were enriched with genes containing TATA-motif promoters, suggesting an association between the depletion of TAF1 and increased expression of genes with TATA-based promoters. Recent structural work in yeast points to an epigenetic role of the TAF1-TAF7 complex in general TFIID function and/or pre-initiation complex (PIC) assembly, and that TAF1 is likely unstable in the absence of a binding partner, such as TAF7 [49].

There is evidence relating various TAFs and the TBP to important functional roles in human neuronal tissue. Indeed, mutations observed in *TAF1-2* and the TBP have been implicated in playing an important role in human neurodegenerative disorders such as X-linked dystonia-parkinsonism (XDP) [59, 60] as well as in intellectual disability and developmental delay [61–63], respectively. Both XDP studies demonstrated aberrant neuron-specific *TAF1* isoform expression levels in neuronal tissue containing *TAF1* mutations. Herzfeld et. al (2013) corroborated previous reports which suggested that a reduction in *TAF1* expression is associated with large-scale expression differences across hundreds of genes. Studies in rat and mice brain also corroborate the importance and relevance of *TAF1* expression patterns specific to neuronal tissues [64, 65]. In corroboration with biochemical and functional studies, a recent population-scale study reported *TAF1* as being ranked 53rd among the top 1,003 constrained human genes [66]. Using allele frequencies reported by the Blood Institute (NHLBI) Exome Sequencing Project (ESP), the authors of this study reasoned that the number of observed missense variants is lower than one would expect by chance (a signed Z score of 5.1779 ranks *TAF1* the 53rd most constrained among the 1,003 most constrained human genes) when compared to the expected probability of missense variants occurring in this gene (5.61E-05).

*FRG2B* and *FAM47B* are not known to be involved in the pathogenesis of human disease, although the detailed molecular function of these genes has been largely unexplored. *FRG2B* is homologous to *FRG2* located on chromosome 4, which has been implicated in playing a role in the pathogenesis of facioscapulohumeral muscular dystrophy (FSHD) in patients with substantial reductions in a 11-150 unit 4q35 microsatellite repeat [67–69]. However, reductions in the homologous 10qt26 microsatellite repeat, proximal to *FRG2B*, have *not* been associated with FSHD.

*ZNF423* acts as a transcriptional regulator and mutations in *ZNF423* coding regions have been implicated in the pathogenesis of Joubert syndrome [70, 71]. The mutation that we have identified in *ZNF423* is located within an intron, and its molecular function is unknown. SLC28A1 is thought to mediate sodiumdepedent fluxes of uridine, adenosine and azidodeoxythymidine [72] whereas SNAPC5, also known as SNAP19, plays a scaffolding role in the forming the complete snRNA-activating protein (SNAP) complex, which is required for the transcription of snRNA genes [73]. The molecular functions of KCHN1 and DMRTB1 are not well understood or studied.

The *TAF1* variant found in this Utah family was the only variant identified as important by the two prioritization schemes that we used. The variant arose in this family as a de novo variant on the X-chromosome of the mother (II-2) of the two affected children (as it is not found in any of the other members of the family) and was then transmitted to both of them. The mother also exhibits extreme X-chromosome skewing, whereas her mother and her sister do not. The protein change occurs within a linker region N-terminal to two bromo domains and C-terminal of a co-factor (TAF7) interacting domain. This linker region is highly conserved in multicellular eukaryotes, but is not present in *S. cerevisiae TAF1*. This linker region is not located in a recently reported crystal structure for the *S. cerevisiae* TAF1-TAF7 complex [49]. The crystal structure of a human TAF1-TAF7 complex was also reported [53], although the characterized region, again, does not include the linker region where our variant lies. This variant represented the only variation that we were able to identify with comprehensive WGS that has a clear molecular function in neuronal tissue and functions as part of TFIID, a larger multi-protein complex involved in general transcription regulation that has been suggested to play a possible role in neurodegenerative diseases and developmental delay when its constituents are disrupted [13, 59–62, 64, 65].

Taken together, this evidence suggests that the variant in *TAF1* may be playing an important role in this newly identified syndrome, whereas other variants found in this family (Table 2) are not as strongly implicated by previous work exploring their putative function(s).

## 5 Conclusions

Our work demonstrates the value of performing more comprehensive genomic analyses when confronted with an undescribed and undiagnosed syndrome or disease affliction, particularly with only one or two affected probands in one family. WGS led to the identification of genomic aberrations that were not tested for by more traditional, clinical assays. Among the multiple rare and potentially disease-contributory variants discovered this this family, the variant in *TAF1* is likely contributing to the phenotype, in concert with the environment and other genetic aberrations to contribute to the sum total of this disease phenotype in the two brothers in Utah. Of course, the differences in genetic background and the environment can certainly account for the phenotypic differences between the two brothers. This phenomenon has been well known for many years in genetics [74, 75], but seems to be more recently appreciated and has become a current active research topic [76–80]. Other work also suggests a physiological link between developmental delay and the TFIID multi-protein complex [13, 61, 62], although the phenotypic variability and expression of other variants in *TAF1* remains to be determined.

## Acknowledgments

G.J.L. is supported by funds from the Stanley Institute for Cognitive Genomics at Cold Spring Harbor Laboratory. K.W. is supported by NIH grant number HG006465. The sequencing by Complete Genomics was provided by a CG data analysis grant to K.W. We would like to thank Max Doerfel for helpful discussions. The authors would like to thank the Exome Aggregation Consortium and the groups that provided exome variant data for comparison. A full list of contributing groups can be found at http://exac.broadinstitute.org/about.

